# Morphological and phylogeographic evidence for budding speciation: an example in hominins

**DOI:** 10.1101/2020.10.23.351114

**Authors:** Caroline Parins-Fukuchi

## Abstract

Parametric phylogenetic approaches that attempt to delineate between distinct ‘modes’ of speciation (splitting cladogenesis, budding cladogenesis, and anagenesis) between fossil taxa have become increasingly popular among comparative biologists. But it is not yet well-understood how clearly morphological data from fossil taxa speak to detailed questions of speciation mode as compared to the lineage diversification models that serve as their basis. In addition, the congruence of inferences made using these approaches with geographic patterns has not been explored. Here, I extend a previously introduced maximum-likelihood approach for the examination of ancestor-descendant relationships to accommodate budding speciation and apply it to a dataset of fossil hominins. I place these results in a phylogeographic context to better understand spatial dynamics underlying the hypothesized speciation patterns. The spatial patterns implied by the phylogeny hint at the complex demographic processes underlying the spread and diversification of hominins throughout the Pleistocene. I also find that inferences of budding are driven primarily by stratigraphic, versus morphological, data and discuss the ramifications for interpretations of speciation process in hominins specifically and from phylogenetic data in general.

## Introduction

Parametric phylogenetic approaches that aim to distinguish between competing ‘modes’ of speciation between fossil taxa have rapidly proliferated over the past several years. These approaches typically employ extended models of lineage diversification that allow taxa to be related by one of several possible speciation patterns that have long been discussed by paleontologists (Stadler et al. 2018). These include: 1) splitting cladogenesis, where a lineage splits evenly into two daughter lineages, 2) budding cladogenesis, where a smaller daughter lineage splits off of an older ancestral lineage, and 3) anagenesis, where a single lineage continuously evolves without splitting. These approaches aim to reconstruct ancestor-descendant (AD) sequences of fossil taxa through the stratigraphic record using a unified model of lineage diversification and stratigraphic preservation (Gavryushkina et al. 2014, Zhang et al. 2016). These approaches have been increasingly leveraged to address empirical patterns in the fossil record (Wright et al. 2020), yielding results that authors have interpreted to highlight the ubiquity of budding speciation as a dominant process in the diversification of new taxa (Bapst and Hopkins 2017). Nevertheless, few investigations have directly examined morphological support for mode. The use of geographic data has also remained under-explored (Fisher 1994, Wright 2017).

Hominins are a strong candidate taxon in which to examine alternative speciation modes in the fossil record, with a history of disagreements over evolutionary mode spanning back decades. Early approaches employed cladograms and stratophenetic diagrams to examine relationships (Delson et al., 1977, Gingerich 1979, Chamberlain and Wood 1987). Repeated attempts to reconstruct hominin phylogeny (Wood 1992, Lieberman et al. 1996, Strait et al. 1997, Strait and Grine 2004, Irish et al. 2013) left major controversies concerning specific hypotheses of direct ancestry unresolved (Strait and Wood 1999, Stringer 2012, Gómez-Robles et al. 2013). Parametric approaches have since been applied to hominins, showing promise in quantifying statistical support for competing hypotheses (Dembo et al. 2015). Further explorations have incorporated AD relationships (Parins-Fukuchi et al. 2019). Despite this improved resolution, many important issues remain in hominin phylogeny. Nevertheless, the role of more complex speciation scenarios such as budding have not been statistically evaluated.

In this study, I examine the morphological and phylogeographic evidence for budding speciation in shaping hominin phylogeny. I extended a previous parametric approach for ‘stratophylogenetic’ inference, including ancestor-descendant relationships, from morphological and stratigraphic data to accommodate budding cladogenesis, which is considered here as a directly ancestral arrangement where a putative ancestor overlaps with a descendant in temporal range. I then use the phylogeny to explore the phylogeographic patterns underlying the spread and morphological divergence of Pleistocene *Homo* populations. Lastly, I evaluate whether morphological data alone support the AD relationships that underpin inferences of budding when the stratigraphic record is not considered.

## Methods and materials

### Data and code availability

All data and tree files used in the analyses are available on Figshare (https://figshare.com/articles/dataset/Morphological_and_phylogeographic_evidence_for_budding_speciation_an_example_in_hominins/13283507). The stratophylogenetic approach was implemented in the Go language as part of the *cophymaru* software package and available as development code on Github (https://github.com/carolinetomo/cophymaru). More detailed information concerning the statistical procedure, including intermediate steps and treatment of the data, is contained within the supplemental text.

### Morphometric data

I used a dataset of cranial landmark coordinates from the literature gathered from 18 fossil hominin specimens and two extant outgroup taxa (González-José et al. 2008). I aligned the 46 3-dimensional landmarks using Procrustes superimposition in MorphoJ (Klingenberg 2011). I then transformed the data using a principal component analysis (PCA) in R (R Core Team 2014). I retained the 13 principal component (PC) axes needed to approach 100% of the variance in the dataset. Phylogenetic inference from morphometric landmarks is viewed as controversial by some (Adams et al. 2011, Varón-González et al. 2020) but appear to perform fairly well in young clades (Caumul and Polly 2005, Goloboff and Catalano 2011, Catalano and Torres 2017, Palci and Lee 2019), such as is the case here.

### Evaluating ancestor-descendant relationships

I extended a previous approach for the evaluation of ancestor-descendant relationships from discrete traits (Parins-Fukuchi et al. 2019) to accommodate both 1) continuous characters and 2) ‘budding’ patterns of ancestry where an ancestral taxon persists after a descendant splits off. Trees were scored using Brownian (Felsenstein 1973) and Poisson (Huelsenbeck and Rannala 1997) models of morphological evolution and stratigraphic preservation, respectively. The implementation developed for this study accepted a fully-bifurcating starting tree as input and identified statistical support for collapsing candidates into direct ancestors. I performed the test using several bifurcating starting trees: the first consistent with the results of Parins-Fukuchi et al. (2019) and successive trials within a nearest-neighbor interchange (NNI) move from the first. Taxa with a lower stratigraphic boundary than their sister taxon or clade were treated as candidate ancestors. I implemented a procedure that iterates through the candidates, collapsing them into ancestors by constraining them to zero branch lengths and assessing statistical support from both the morphological and stratigraphic data. I retained the topology, including AD relationships, from the tree with the best Akaike information criterion (AIC) score. AIC was used rather than raw log-likelihoods because the addition of an AD relationship reduces the number of free parameters by one. Model uncertainty for AD assignments was estimated using AIC weights (Wegenmakers and Farrell 2004). All AD analyses were performed using the *mandos* executable available in the *cophymaru* code repository (cited above).

### Biogeographic analyses

I inferred biogeographic ranges using Lagrange (https://github.com/blackrim/lagrange). Internal nodes associated with ancestral taxa were constrained to geographic ranges corresponding to the associated taxon. Ranges were treated as discrete units corresponding to four areas occupied by hominin populations throughout the Plio-Pleistocene: Africa, Asia, the Middle East (encompassing the Levant, the Arabian Peninsula, and the Steppes), and Europe.

## Results and Discussion

### Budding speciation in hominin evolution

When considered in a stratophylogenetic framework, morphometric and temporal data suggest several possible episodes of budding speciation (Figs. 1, S2). Neither of the *Australopithecus* samples included in the analysis were identified as ancestral taxa. Like several previous studies, *A. afarensis* is recovered as an outgroup to a clade comprising all of the later-occurring hominin morphotaxa. Many authors have interpreted those cladistic results to imply that *A. afarensis* is ancestral to *Homo*. However, these results suggest that *A. afarensis* is not directly ancestral but is instead a terminal outgroup. Since *A. anamensis* appears to be a chronospecies that is ancestral to *A. afarensis* (Kimbel 2006, Parins-Fukuchi et al. 2019), it is possible that the specimens assigned to *A. anamensis* represent a population ancestral to both *A. afarensis* and the later-occurring hominin lineages. *P. boisei* is identified as the direct ancestor of *P. robustus*. This finding is consistent with previous qualitative interpretations (Kimbel 2007). *P. aethiopicus*, on the other hand, is reconstructed with high confidence to be a distinct, branching taxon (Table 1).

**Figure 1:**
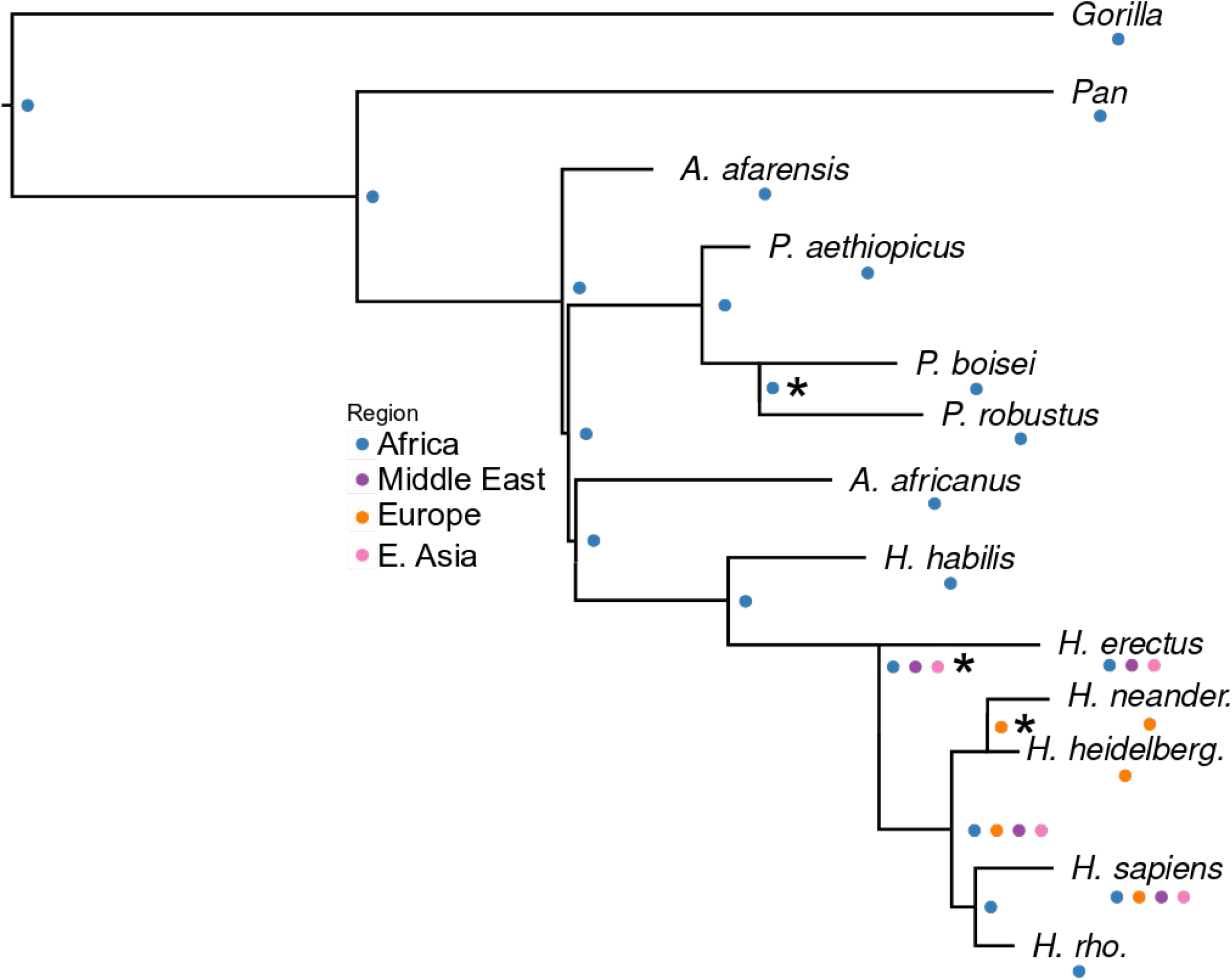
Best stratophylogenetic tree with geographic range reconstructions mapped to internal nodes. Asterisks denote nodes fixed in the biogeographic analysis due to their reconstruction as ancestral taxa. Geographic ranges representing tips and sampled ancestral taxa correspond only to the particular specimens contributing to the morphological data within each taxon in this study (see table 2 for full list of specimens).

**Table 1.** Support for each candidate taxon’s position as a direct ancestor on the best stratophylogenetic tree. Model support for each directly ancestral assignment is given by the AIC weight of the ancestral arrangement calculated relative to the model support for the bifurcating arrangement (see methods). The resulting support indices fall between 0-1, with values above 0.5 indicating greater support for a budding or anagenetic arrangement and below 0.5 indicating preference for a bifurcating relationship. When calculating support for each taxon, all other relationships and AD assignments in the tree were held at their ML placements. The first column reflects AIC support yielded by the combined stratigraphic and morphological data set, while the second column reflects support displayed by morphology alone.

| OTU | Combined AIC weight | Morphological AIC weight |
| --- | --- | --- |
| *P. aethiopicus* | 0.03 | 0.02 |
| *P. boisei* | 0.75 | 0.51 |
| *A. afarensis* | 0.02 | 0.02 |
| *A. africanus* | 0.14 | 0.02 |
| *H. habilis* | 0.41 | 0.21 |
| *H. erectus* | 0.77 | 0.26 |
| *H. heidelbergensis* | 0.75 | 0.49 |
| *H. rhodesiensis* | 0.02 | 0.01 |

The analysis revealed support for a budding event between *H. erectus* and the clade encompassing *H. heidelbergensis, H. rhodesiensis*, Neanderthals, and modern humans. The mid-Pleistocene specimens corresponding to *H. heidelbergensis* and *H. rhodesiensis* grouped polyphyletically, forming clades with Neanderthals and *H. sapiens*, respectively. This result is notable given the historical tendency to treat specimens corresponding to each taxon (as treated here) as geographic variants of *H. heidelbergensis. H. heidelbergensis*, represented by only European specimens in this study, was reconstructed as a budding ancestor to Neanderthals, while *H. rhodesiensis* was reconstructed as a sister taxon to modern humans. The placement of European *H. heidelbergensis* specimens as directly ancestral to Neanderthals is consistent with both the genomic results of Meyer et al. (2016) and the morphological results of Mounier and Caparrós (2016), the former of which inferred the Atapuerca specimens as most closely related to Neanderthals and the latter of which reconstructed both Atapuerca 5 and Steinheim (the two specimens representing *H. heidelbergensis* in this study) as outgroups to Neanderthals. It does however, disagree with a morphospecies-level phylogenetic analysis, which inferred *H. heidelbergensis*, defined as including *both* the European and African samples included here, as ancestral to both Neanderthals and modern humans (Parins-Fukuchi et al. 2019). The discrepancy between studies can therefore be explained most simply by their differing treatments of *H. heidelbergensis*. The results here further demonstrate that careful examination of geography and morphology below the morphospecies level are needed to develop a thorough understanding of the spatial and demographic processes that shaped the divergence and reticulation of hominin populations throughout the Pleistocene.

Support of AD relationships differs when temporal and morphological data are considered together and when morphology is considered separately (Table 1, Fig. S3-S4). In contrast to the combined dataset, the morphological data alone are either equivocal (*H. heidelbergensis*, *P. boisei*) or prefer a bifurcating arrangement (*H. erectus*). This may reflect the limits of morphospecies-level data. The temporal and spatial heterogeneity expected in a long-lived and widespread lineage such as *H. erectus* (Baab 2008) may simply yield too much demographic complexity to map neatly to such a coarse representation of process. Adopting a finer scale of analysis will facilitate testing of the phylogenetic cohesiveness of named Pleistocene taxa such as *H. erectus* and *H. heidelbergensis* and identify whether breaking down spatial heterogeneity can yield a more detailed understanding of the processes underlying the divergence of Pleistocene populations.

### Pleistocene biogeography

The biogeographic range reconstructions suggest the presence of a geographically widespread, mid-Pleistocene ancestor to humans, Neanderthals, *H. heidelbergensis*, and *H. rhodesiensis*. Pleistocene *Homo* can thus be characterized by patterns in widespread dispersal, followed by gradual fragmentation into geographically-distinct subpopulations. *H. heidelbergensis* and *H. rhodesiensis* are separated into geographically separate clades, disagreeing with treatments lumping them together into a single geographically-widespread *H. heidelbergensis*. These results further underscore the need for examination at lower taxonomic scales than is typically undertaken. Overall, the picture of hominin evolution presented here represents an alignment of phylogenetic results with the heterogeneous geographic patterns observed in *H. erectus* and early African *H. sapiens* (Baab 2011, Hublin et al. 2017) and the complicated network of genetic interactions (Green et al. 2010, Kuhlwilm et al. 2016, Villanea and Schraiber 2019, Rogers et al. 2020) toward a deeper statistical understanding of the complexity in evolutionary and demographic patterns between hominin populations throughout the Pleistocene.

### Phylogenetics and speciation processes in the fossil record

This study highlights the difficulty in identifying ancestral fossil populations from taxonomically-coarse data. The comparatively weak morphological support for the AD relationships identified here suggests that researchers should be cautious when interpreting paleontological phylogenies. In general, I suggest that reconstructions of AD relationships that are generated from phylogenetic data should generally be treated as contingent and coarse reconstructions as opposed to detailed inferences of the processes underlying the divergence of species. Population-level phenotypic (or molecular) and spatial data are typically needed to generate a detailed understanding of speciation processes in living and fossil taxa. If satisfied with the particular scheme of OTU resolution and a coarse scale of inference, the framework here may be broadly useful to discern general patterns in phylogenetic ancestry. However, researchers using approaches that seek to delineate between competing modes of speciation should be mindful of scale. Further development of a clear understanding of the capabilities and limitations of data at multiple timescales will enable a continued expansion of the boundaries of fossil data when speaking to fundamental questions in evolutionary biology.

## Acknowledgements

I thank L MacLatchy, Z Alemseged, DC Fisher, M Foote, SA Smith, G Auteri, J Saulsbury, and GW Stull for discussions that greatly benefited this work. The author was supported as a TC Chamberlin Postdoctoral Scholar in the Department of Geophysical Sciences at the University of Chicago while undertaking and completing this work.

## References

Baab, Karen L. “The taxonomic implications of cranial shape variation in Homo erectus.” Journal of Human Evolution 54.6 (2008): 827–847.

Baab, Karen L. “Cranial shape in Asian Homo erectus: geographic, anagenetic, and size-related variation.” In Asian paleoanthropology, C.J. Norton and D.R. Braun, eds. Springer, Dordrecht, (2011): 57–79.

Bapst, David W., and Melanie J. Hopkins. “Comparing cal3 and other a posteriori time-scaling approaches in a case study with the pterocephaliid trilobites.” Paleobiology 43.1 (2017): 49–67.

Chamberlain, A. T., and Bernard A. Wood. “Early hominid phylogeny.” Journal of Human Evolution 16.1 (1987): 119–133.

Delson, Eric, Niles Eldredge, and Ian Tattersall. “Reconstruction of hominid phylogeny: a testable framework based on cladistic analysis.” Journal of Human Evolution 6.3 (1977): 263–278.

Dembo, Mana, Nicholas J. Matzke, Arne Ø. Mooers, and Mark Collard. “Bayesian analysis of a morphological supermatrix sheds light on controversial fossil hominin relationships.” Proceedings of the Royal Society B: Biological Sciences 282.1812 (2015): 20150943.

Felsenstein, Joseph. “Maximum-likelihood estimation of evolutionary trees from continuous characters.” American journal of human genetics 25.5 (1973): 471.

Felsenstein, Joseph. “Quantitative characters, phylogenies, and morphometrics.” in Morphology, Shape and Phylogeny 64 (2002).

Fisher, Daniel C. “Stratocladistics: integrating temporal data and character data in phylogenetic inference.” Annual Review of Ecology, Evolution, and Systematics 39 (2008): 365–385.

Fisher, Daniel C. “Stratocladistics: morphological and temporal patterns and their relation to phylogenetic process.” In: Interpreting the hierarchy of nature. Grande, and O. Rieppel, eds. (1994): 133–171.

Foley, Robert. “Species diversity in human evolution: challenges and opportunities.” Transactions of the Royal Society of South Africa 60.2 (2005): 67–72.

Gavryushkina, Alexandra, David Welch, Tanja Stadler, and Alexei J. Drummond. “Bayesian inference of sampled ancestor trees for epidemiology and fossil calibration.” PLoS Comput Biol 10, no. 12 (2014): e1003919.

Gingerich, Philip D. “The stratophenetic approach to phylogeny reconstruction in vertebrate paleontology.” In Phylogenetic analysis and paleontology, J. Cracraft, and N. Eldredge, eds. (1979): 41–77.

Gómez-Robles, Aida, José María Bermúdez de Castro, Juan-Luis Arsuaga, Eudald Carbonell, and P. David Polly. “No known hominin species matches the expected dental morphology of the last common ancestor of Neanderthals and modern humans.” Proceedings of the National Academy of Sciences 110.45 (2013): 18196–18201.

González-José, Rolando, Rolando, Ignacio Escapa, Walter A. Neves, Rubén Cúneo, and Héctor M. Pucciarelli. “Cladistic analysis of continuous modularized traits provides phylogenetic signals in Homo evolution.” Nature 453.7196 (2008): 775.

Gould, Stephen Jay, and Niles Eldredge. “Punctuated equilibria: the tempo and mode of evolution reconsidered.” Paleobiology 3.2 (1977): 115–151.

Green, Richard E., Johannes Krause, Adrian W. Briggs, Tomislav Maricic, Udo Stenzel, Martin Kircher, Nick Patterson et al. “A draft sequence of the Neandertal genome.” Science 328.5979 (2010): 710–722.

Hublin, Jean-Jacques, Ben-Ncer, A., Bailey, S.E., Freidline, S.E., Neubauer, S., Skinner, M.M., Bergmann, I., Le Cabec, A., Benazzi, S., Harvati, K. and Gunz, P. “New fossils from Jebel Irhoud, Morocco and the pan-African origin of Homo sapiens.” Nature 546.7657 (2017): 289.

Huelsenbeck, John P., and Bruce Rannala. “Maximum likelihood estimation of phylogeny using stratigraphic data.” Paleobiology 23.2 (1997): 174–180.

Johanson, Donald C., and Tim D. White. “A systematic assessment of early African hominids.” Science 203.4378 (1979): 321–330.

Kimbel, William H. The Species and Diversity of Australopiths. In: Handbook of Paleoanthropology. Springer, Berlin. (2007)

Kimbel, William H., et al. “Was Australopithecus anamensis ancestral to A. afarensis? A case of anagenesis in the hominin fossil record.” Journal of Human Evolution 51.2 (2006): 134–152.

Klein, Richard G. “Darwin and the recent African origin of modern humans.” (2009): 16007–16009.

Klingenberg, Christian Peter. “MorphoJ: an integrated software package for geometric morphometrics.” Molecular ecology resources 11.2 (2011): 353–357.

Kuhlwilm, Martin, Ilan Gronau, Melissa J. Hubisz, Cesare De Filippo, Javier Prado-Martinez, Martin Kircher, Qiaomei Fu et al. “Ancient gene flow from early modern humans into Eastern Neanderthals.” Nature 530.7591 (2016): 429–433.

Lieberman, Daniel E., Bernard A. Wood, and David R. Pilbeam. “Homoplasy and early Homo: an analysis of the evolutionary relationships of H. Habilis sensu stricto and H. rudolfensis.” Journal of Human Evolution 30.2 (1996): 97–120.

Meyer, Matthias, Juan-Luis Arsuaga, Cesare de Filippo, Sarah Nagel, Ayinuer Aximu-Petri, Birgit Nickel, Ignacio Martínez et al. “Nuclear DNA sequences from the Middle Pleistocene Sima de los Huesos hominins.” Nature 531, no. 7595 (2016): 504–507.

Mounier, A., and Miguel Caparrós. “The phylogenetic status of Homo heidelbergensis–a cladistic study of Middle Pleistocene hominins.” Bmsap 27.3-4 (2015): 110–134.

Parins-Fukuchi, Caroline. “Bayesian placement of fossils on phylogenies using quantitative morphometric data.” Evolution 72.9 (2018): 1801–1814.

Parins-Fukuchi, Caroline, Elliot Greiner, Laura M. MacLatchy, and Daniel C. Fisher. “Phylogeny, ancestors, and anagenesis in the hominin fossil record.” Paleobiology 45.2 (2019): 378–393.

R Core Team (2014). R: A language and environment for statistical computing. R Foundation for Statistical Computing, Vienna, Austria. URL http://www.R-project.org/

Rogers, Alan R., Nathan S. Harris, and Alan A. Achenbach. “Neanderthal-Denisovan ancestors interbred with a distantly related hominin.” Science advances 6.8 (2020): eaay5483.

Silvestro, Daniele, Rachel CM Warnock, Alexandra Gavryushkina, and Tanja Stadler. “Closing the gap between palaeontological and neontological speciation and extinction rate estimates.” Nature Communications 9, no. 1 (2018): 1–14.

Strait, David S., and Frederick E. Grine. “Inferring hominoid and early hominid phylogeny using craniodental characters: the role of fossil taxa.” Journal of Human Evolution 47.6 (2004): 399–452.

Strait, David S., Frederick E. Grine, and Marc A. Moniz. “A reappraisal of early hominid phylogeny.” Journal of Human Evolution 32.1 (1997): 17–82.

Strait, David S., and Bernard A. Wood. “Early hominid biogeography.” Proceedings of the National Academy of Sciences 96.16 (1999): 9196–9200.

Stringer, Chris B. “The status of Homo heidelbergensis (Schoetensack 1908).” Evolutionary Anthropology: Issues, News, and Reviews 21.3 (2012): 101–107.

Stadler, Tanja, Alexandra Gavryushkina, Rachel CM Warnock, Alexei J. Drummond, and Tracy A. Heath. “The fossilized birth-death model for the analysis of stratigraphic range data under different speciation modes.” Journal of theoretical biology 447 (2018): 41–55.

Villanea, Fernando A., and Joshua G. Schraiber. “Multiple episodes of interbreeding between Neanderthal and modern humans.” Nature Ecology & Evolution 3.1 (2019): 39–44.

Wagenmakers, Eric-Jan, and Simon Farrell. “AIC model selection using Akaike weights.” Psychonomic Bulletin and Review 11.1 (2004): 192–196.

Wood, Bernard. “Early hominid species and speciation.” Journal of Human Evolution 22.4-5 (1992): 351–365.

Wright, April, Peter Wagner, and David Wright. “Testing character-evolution models in phylogenetic paleobiology: a case study with Cambrian echinoderms.” (2020).

Wright, David F. “Bayesian estimation of fossil phylogenies and the evolution of early to middle Paleozoic crinoids (Echinodermata).” Journal of Paleontology 91.4 (2017): 799–814.

Zhang, Chi, Tanja Stadler, Seraina Klopfstein, Tracy A. Heath, and Fredrik Ronquist. “Total-evidence dating under the fossilized birth–death process.” Systematic biology 65, no. 2 (2016): 228–249.

