## Supplemental methods for "Morphological and phylogeographic evidence for budding speciation: an example in hominins"

*Stratigraphic range data*: Stratigraphic ranges for hominin taxa were taken from Parins-Fukuchi et al. (2019a, 2019b).

*Treatment of morphometric data*: I transformed the landmark data into statistically independent variables using principal components analysis (PCA). The study from which the landmark data were obtained (González-José et al. 2008) also transformed the data using PCA, however, they did so by manually dividing the landmarks into modules *a priori* and performing PCA on each module, retaining only the first axis. In this study, I pooled the landmarks (after Goloboff and Catalano 2011), ignoring the authors' original 'modularized' treatment. This is because the modules defined by the authors were not tested statistically and so their biological meaningfulness is unclear-- a major criticism of the study (Adams et al. 2011). I retained the PC axes from the pooled sample that asymptotically approached 100% of the variance in the dataset. The fact that each axis describes a different proportion of the overall variance was taken into account naturally because the ranked nature of the axes causes each component to contribute differently to the calculation of the morphological branch lengths, with the most variable (PC1) contributing more than the least (PC13). Nevertheless, not necessarily clear what the most appropriate handling would be, given that many phylogeneticists have argued to hold all characters on equal footing in terms of how their evidence is weighted. While worthwhile for future investigations, this theoretical discussion falls outside the scope of this particular study.

Ideally, future methodological extensions would treat morphometric data without PCA transformation under an approach that iteratively optimizes the landmark alignment and the phylogenetic topology, such as is done using the parsimony-based 'phylogenetic morphometrics' approach introduced by Catalano and Goloboff (2010). Nevertheless, inference from PC scores under maximum-likelihood likely perform adequately well (Caumul and Polly 2005) for the synthetic approach used here. While they perform better than PC scores (Catalano and Torres 2017), more sophisticated treatments of landmark data impart only a slight improvement. In addition, an approach akin to Catalano and Goloboff's has not yet been implemented in a parametric context, despite being achievable in principle. Nevertheless, although imperfect, the qualitatively reasonable performance of geometric morphometric data over fairly short timescales (such as is employed here) support their use here. In addition, the quantitative treatment used here, even if imperfect, has the advantage of reducing uncertainty stemming from subjective and conflicting accounts of discrete character states that may have hampered reconciliation between previous studies using traditional cladistic methods. Thus, the trade-offs are likely appropriate for the examination undertaken here.

*Assessment of ancestor-descendant relationships*: The general methodology by which I evaluated ancestor-descendant (AD) relationships in this study was based on that employed by Parins-Fukuchi et al. (2019a). Their approach involved the testing of ancestor-descendant relationships between sister lineages with non-overlapping temporal ranges using discrete character and stratigraphic ranges expressed as intervals of continuous time. As a result, the previous approach only considered ancestor-descendant relationships between serially-linked taxa, facilitating the examination of only anagenetic and bifurcating (splitting cladogenetic) relationships. I extended that previous approach in two ways. 1) I implemented a Brownian motion (BM) model of trait evolution to accommodate continuous traits within the previously-developed framework and 2) I enabled the examination of AD relationships between taxa with overlapping temporal ranges, thus extending the set of considered speciation scenarios to include budding cladogenesis.

Like Parins-Fukuchi et al., stratigraphic likelihoods were calculated using a Poisson model of fossil preservation. This model is a homogeneous Poisson process, with the rate parameter corresponding to the rate of fossil deposition. Readers can consult Huelsenbeck and Rannala (1997) for a full derivation of the stratigraphic likelihood function or Parins-Fukuchi et al. (2019a) for a more concise mathematical treatment. As implemented in a phylogenetic context, the model favors arrangements that minimize the amount of unsampled 'ghost lineage' time implied by the phylogeny. As a result, when given a pair of taxa with different lower boundaries in the stratigraphic record, the stratigraphic likelihood will always improve when making a bifurcating split into an AD arrangement (Fig. S1). AD relationships were also evaluated under a Brownian model of morphological evolution. BM likelihoods were calculated along the phylogeny with branch lengths proportional to units of Gaussian variance which were optimized using an expectation-maximation algorithm after Felsenstein (1981) and implemented by Parins-Fukuchi (2018). Trees were thus evaluated under two different sets of branch lengths: proportional to time for stratigraphic data and to morphological disparity for the character data. This was also the case in the discrete-character-based approach of Parins-Fukuchi et al. (2019a). The best-scoring topology with Brownian branch lengths is presented in Figure S2. Taxa were made into ancestors by fixing their branch lengths to zero. AD relationships were tested by comparing Akaike Information Criterion (AIC) values based upon summed morphological and stratigraphic likelihoods between bifurcating and AD arrangements.

The implementation developed for this study is contained within the *mandos* executable hosted on the development branch of the *cophymaru* repository on Github (https://github.com/carolinetomo/cophymaru) at the time of this writing. The approach examines AD relationships in a semi-automated manner. From a user-supplied starting topology, the procedure iteratively collapses and expands terminal taxa into direct ancestors. Only those taxa with stratigraphic ranges older than those of their sister taxon or clade were considered. I manually generated a starting tree that was congruent with the results of Parins-Fukuchi et al. (2019a), which represented a superset of the taxa examined here. I then created several alternative bifurcating topologies that were within a nearest-neighbor interchange (NNI) move from the initial tree. I then used the *mandos* executable to identify the best-supported AD arrangement from each starting topology and retained the AD tree with the lowest AIC score. Future extensions will automate the NNI rearrangements to the starting tree, but the manual rearrangements done here were adequate given the small number of taxa present. The AD trees can be expressed with branch lengths either proportional to time, representing the stratigraphic record, or morphological divergence, as calculated from the morphometric data (with direct ancestors displaying branch lengths of zero). Final AD trees with both stratigraphic and morphological branch lengths calculated for all of the tested bifurcating topologies are available in the data supplement (https://figshare.com/articles/dataset/Morphological_and_phylogeographic_evidence_for_budding_speciation_an_example_in_hominins/13283507 – cited in main text).

**Supplemental figures:**

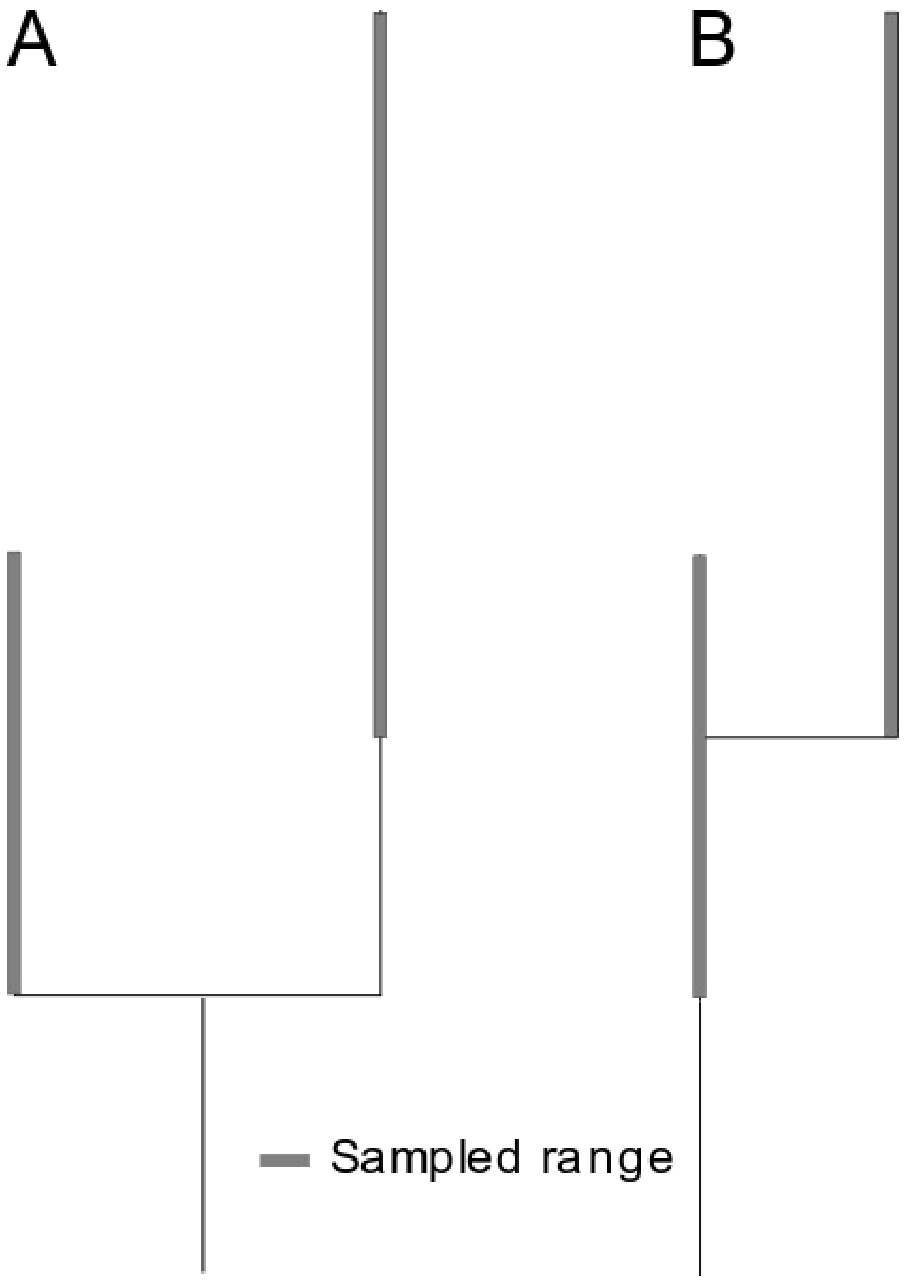


*Supplemental figure 1*. Transformation of bifurcating phylogenetic relationship where sister taxa are descended from a hypothetical common ancestor to a budding relationship where the earlier-occurring taxon represents the direct ancestor of the later-occurring taxon. The budding arrangement reduces the amount of unsampled evolutionary time implied by the phylogeny.

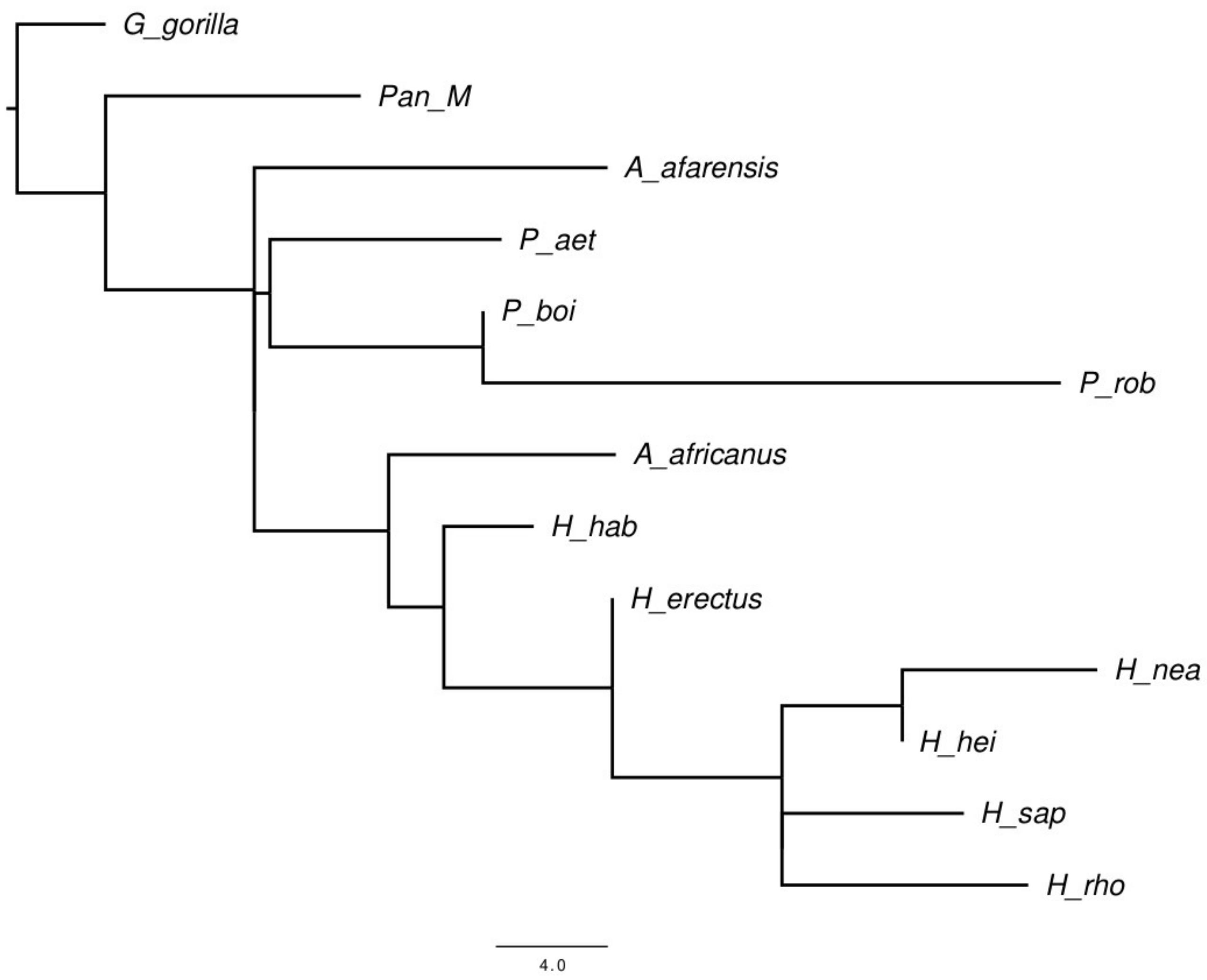


*Supplemental figure 2*. Best scoring topology (including AD arrangements) with branch lengths proportional to morphological disparity expressed in units of Brownian variance. Direct ancestors are represented as taxa with zero branch lengths.

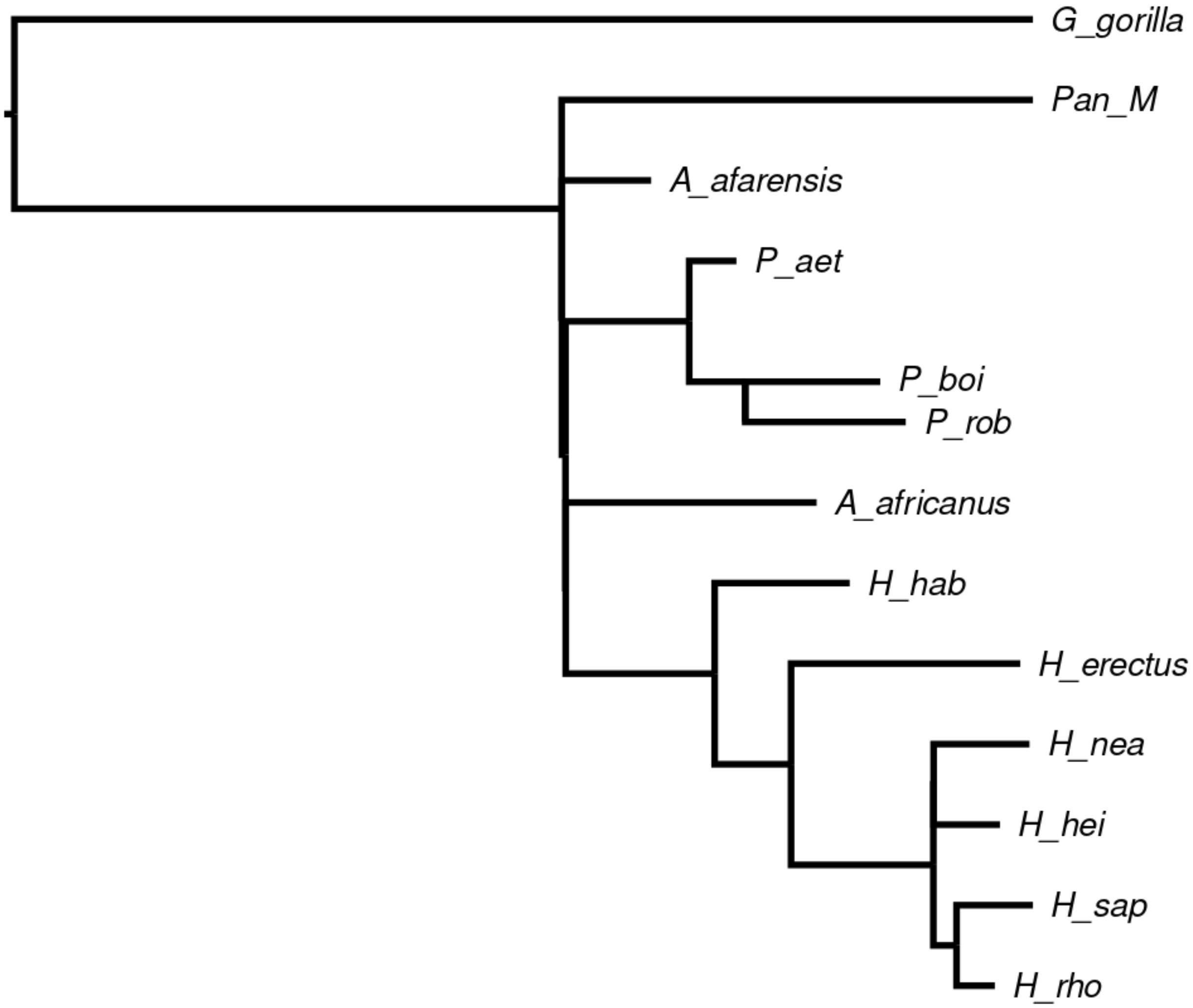


*Supplemental figure 3*. Best-scoring tree with only the direct ancestors that are supported morphological data alone. The starting topology is thus the same as is presented in Fig. 1, but only *P. boisei* is retained as an ancestor, given the lack of morphological support for the other ancestors preferred when morphology and stratigraphy are considered together.

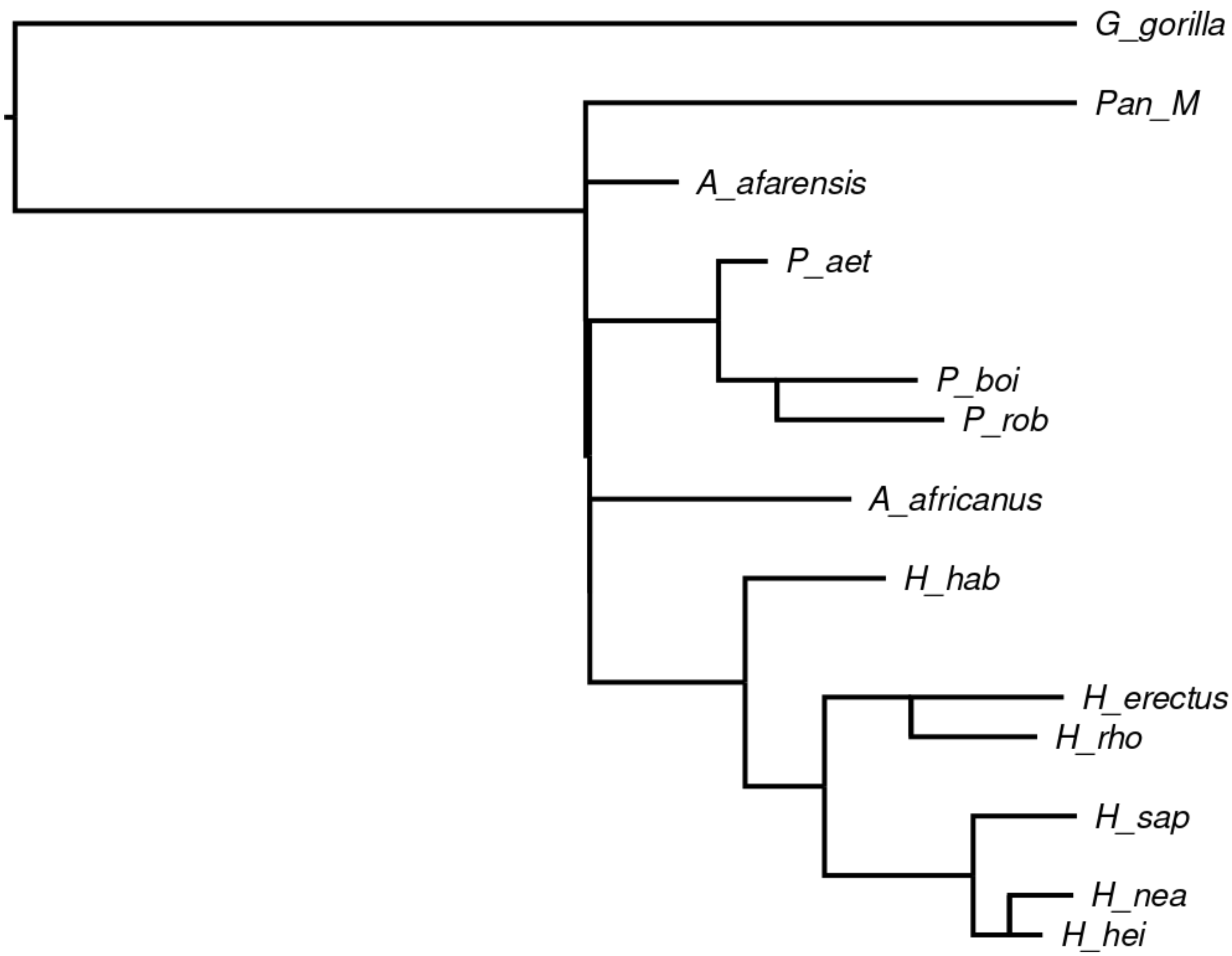


*Supplemental figure 4*. Best-scoring tree overall when only morphological data are considered. This topology differs from the tree recovered from the combined dataset (presented in Fig. 1) by one nearest-neighbor interchange move and several ancestors.

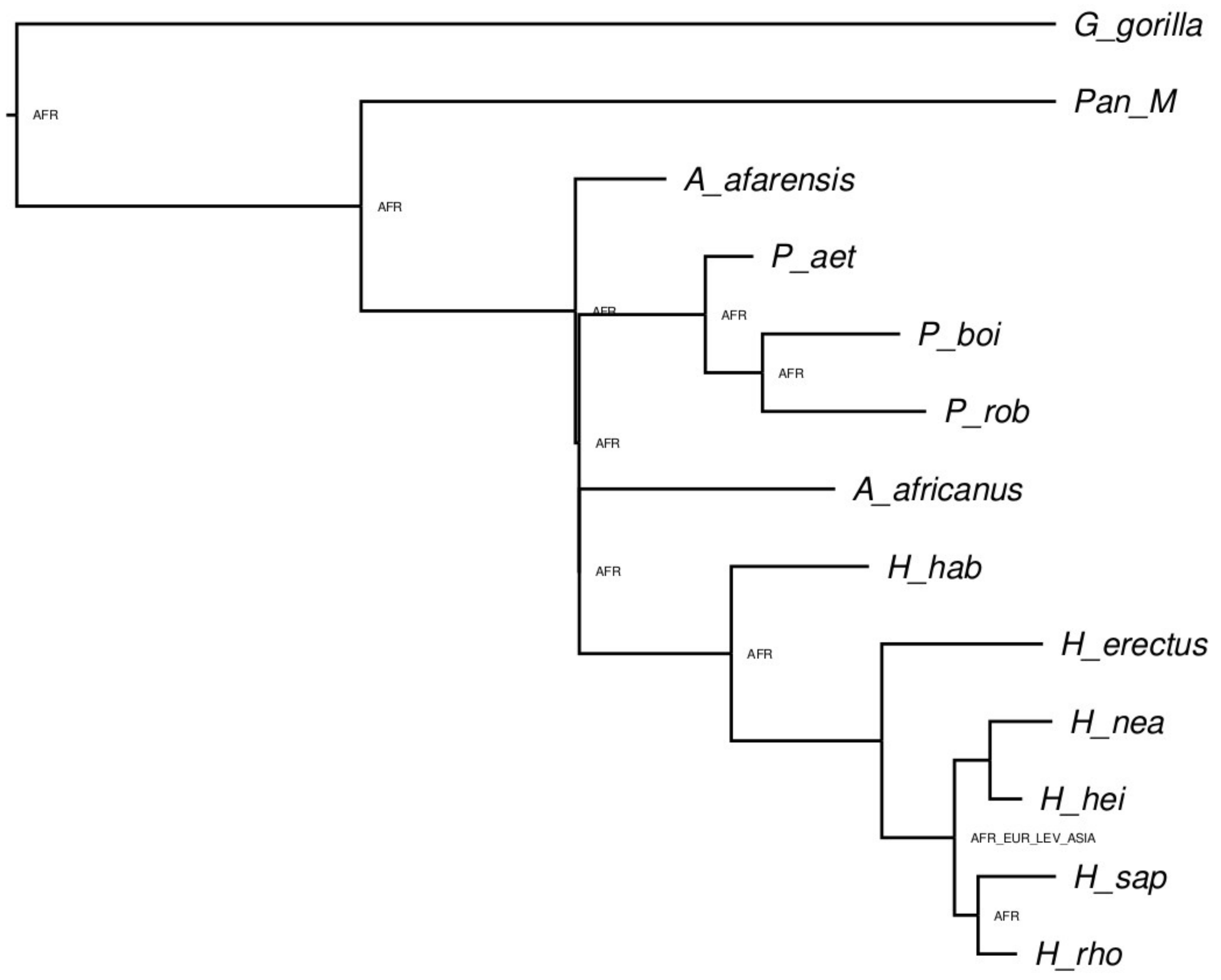


*Supplemental figure 5*. Raw lagrange output. Internal nodes without marked ranges correspond to the direct ancestors retained in the tree presented in Fig. 1. They were fixed in the analysis to reflect the ranges displayed in the fossil record. The full configuration file, including constrained ancestral ranges, can be found in the data supplement.

**Supplemental Tables:**

| Specimen | OTU | Label |
|---|---|---|
| A.L.444-2 | *Australopithecus afarensis* | A_afarensis |
| Sts 5 | *Au. africanus* | A_africanus |
| KNMER-406 | *Paranthropus boisei* | P_boi_406 |
| OH 5 | *P. boisei* | P_boi_OH5 |
| SK 48 | *P. robustus* | P_rob |
| WT 17000 | *P. aethiopicus* | P_aet |
| KNMER 1470 | *Homo habilis* | P_rud |
| KNMER 1813 | *H. habilis* | H_hab |
| KNMER 3733 | *H. erectus* | H_erg_1 |
| D2700 | *H. erectus* | H_erg_2 |
| Zhoukoudian | *H. erectus* | H_ere |
| Steinheim | *H. heidelbergensis* | H_hei_S |
| Broken Hill 1 | *H. heidelbergensis* | H_hei_BH |
| Atapuerca 5 | *H. heidelbergensis* | H_hei_A |
| Gibraltar 1, Forbes' Quarry | *H. neanderthalensis* | H_nea_G |
| La Chappelle-au Saints 1 | *H.neanderthalensis* | H_nea_LC |
| La Ferrassie 1 | *H. neanderthalensis* | H_nea_LF |
| Patagonian, Rio Negro #797 | *H. sapiens* | H_sap |

*Table S1*: Specimens used in the morphometric dataset. Additional details concerning the museum storage of specimens are available from the original study (González-José et al. 2008).

**Supplemental References:**

Adams, D.C., Cardini, A., Monteiro, L.R., O'Higgins, P. and Rohlf, F.J. “Morphometrics and phylogenetics: principal components of shape from cranial modules are neither appropriate nor effective cladistic characters.” Journal of Human Evolution, 60.2 (2011): 240-243.

Caumul, Radhekshmi, and P. David Polly. "Phylogenetic and environmental components of morphological variation: skull, mandible, and molar shape in marmots (Marmota, Rodentia)." Evolution 59.11 (2005): 2460-2472.

Catalano, Santiago A., and Ambrosio Torres. “Phylogenetic inference based on landmark data in 41 empirical data sets.” Zoologica Scripta 46.1 (2017): 1-11.

Catalano, Santiago A., Pablo A. Goloboff, and Norberto P. Giannini. "Phylogenetic morphometrics (I): the use of landmark data in a phylogenetic framework." Cladistics 26.5 (2010): 539-549.

Felsenstein, Joseph. "Evolutionary trees from gene frequencies and quantitative characters: finding maximum likelihood estimates." Evolution (1981): 1229-1242.

Goloboff, Pablo A., and Santiago A. Catalano. "Phylogenetic morphometrics (II): algorithms for landmark optimization." Cladistics 27.1 (2011): 42-51.

González-José, Rolando, Rolando, Ignacio Escapa, Walter A. Neves, Rubén Cúneo, and Héctor M. Pucciarelli. "Cladistic analysis of continuous modularized traits provides phylogenetic signals in Homo evolution." *Nature* 453.7196 (2008): 775.

Huelsenbeck, John P., and Bruce Rannala. "Maximum likelihood estimation of phylogeny using stratigraphic data." Paleobiology (1997): 174-180.s

Palci, Alessandro, and Michael SY Lee. "Geometric morphometrics, homology and cladistics: review and recommendations." Cladistics 35.2 (2019): 230-242.

Parins-Fukuchi, Caroline. "Bayesian placement of fossils on phylogenies using quantitative morphometric data." Evolution 72.9 (2018): 1801-1814.

Parins-Fukuchi, Caroline, Elliot Greiner, Laura M. MacLatchy, and Daniel C. Fisher. "Phylogeny, ancestors, and anagenesis in the hominin fossil record." *Paleobiology* 45.2 (2019a): 378-393.

Parins-Fukuchi, Caroline, Elliot Greiner, Laura M. MacLatchy, and Daniel C. Fisher. Data from: Phylogeny, ancestors, and anagenesis in the hominin fossil record, Dryad, Dataset. (2019b): https://doi.org/10.5061/dryad.6d2h8bp
